# Facile determination of the Poisson’s ratio and Young’s modulus of polyacrylamide gels and polydimethylsiloxane

**DOI:** 10.1101/2023.05.10.540222

**Authors:** Ariell M. Smith, Dominique Gabriele Inocencio, Brandon M. Pardi, Arvind Gopinath, Roberto Andresen Eguiluz

## Abstract

Polyacrylamide hydrogels (PAH) and polydimethylsiloxane (PDMS) are two soft materials often used in cell mechanics and mechanobiology, in manufacturing lab-on-a chip applications, among others. This is partly due to the ability to tune their elasticity with ease, in addition to various chemical modifications. For affine polymeric networks, two (of three) elastic constants – the Young’s modulus (*E*), the shear modulus (*G*), and the Poisson’s ratio (*ν*) – describe the purely elastic response to external forces. However, the literature addressing the experimental determination of ν for PAH (also sometimes referred to as PAA gels in the literature) and PDMS is surprisingly limited when compared to the literature reporting values of *E* and *G*. Here, we present a facile method to obtain the Poison’s ratio and Young’s modulus for PAH and PDMS based on static tensile tests, and cross-correlate these values with those obtained via a second independent method, shear rheology. We show that: i) the Poisson’s ratio may vary significantly from the value for incompressible materials (ν = 0.5), and ii) find a high degree of agreement between shear rheology and macroscopic static tension tests for PAH but not PDMS.

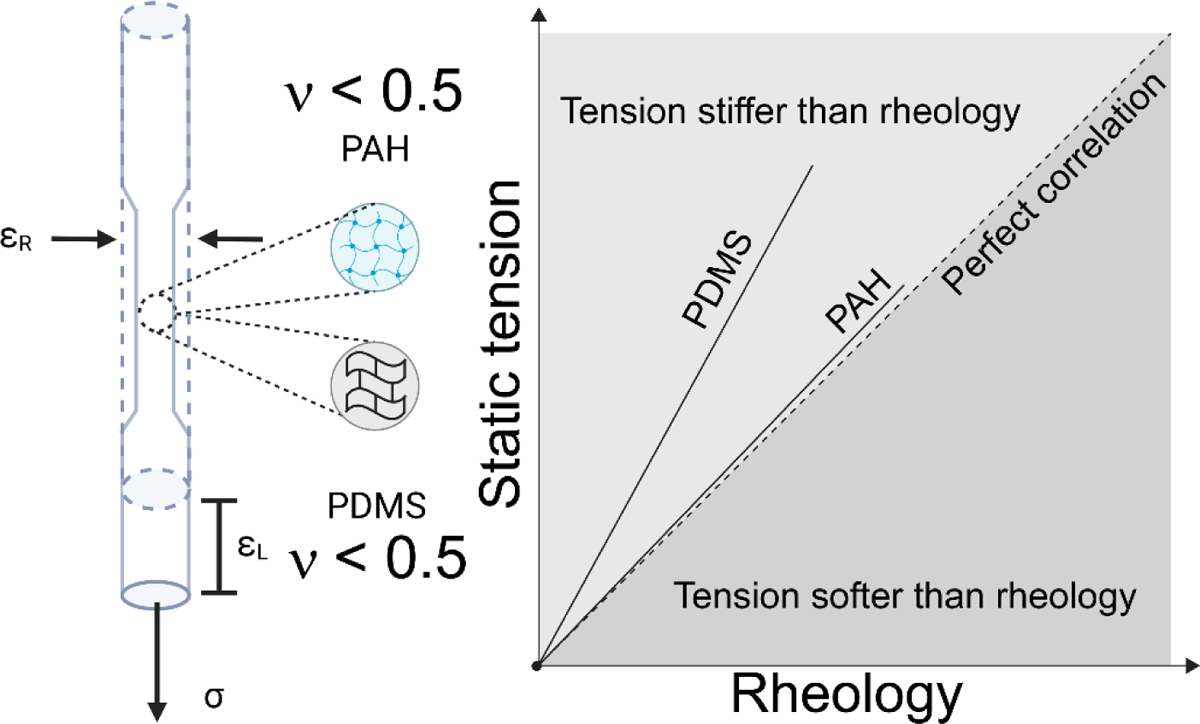

## INTRODUCTION

Gels and elastomers have become popular substrates for studies of cell mechanics and mechanobiology, in manufacturing lab-on-a chip applications, among others.^1–5^ This is in part due to the ease in fabricating these materials into complex shapes, the ability to tune the mechanical properties and the variety of specifically tailored surface modifications possible. These soft materials may be characterized mechanically at a variety of scales: elastic properties can be determined at small scales using atomic force microscopy (AFM), at larger bulk scales by shear and normal rheology, or by dynamic mechanical analyses using indentation or tension. Since these materials possess strain and strain-rate dependent properties and are often hydrated and porous, rheology provides an especially efficient mode of interrogation and analysis.

For small deformations and under the action of weak stresses, gels and elastomers behave as linear elastic materials. When coarse-grained and probed at the macroscale, these materials may be also approximated as isotropic materials. Linear isotropic elastic materials are characterized by three moduli characterizing different deformation modes - the extensional modulus *K* (addressing extensional properties), the shear modulus *G* (addressing shearing deformations), and the bulk modulus *B* (quantifying response to bulk volumetric compression). These elastic moduli depend linearly on the Young’s modulus *E*, and are related by the Poisson’s ratio ν^5–12^ that quantifies the compressibility of the material and is therefore an important material property for soft gels. While *K* and *G* are both measures of the stiffness (or how much the material resists change in shape), ν describes the coupling between axial and transverse deformations. That is, the Poisson’s ratio ν quantifies the degree to which the material contracts laterally (under axial tension) or expands laterally (under axial compression) under applied loads.^13^ Ideal incompressible materials such as rubber, maintain their volume under load (corresponding to high values of *B*) and only change their shape, with ν = 0.5. Soft hydrogels and many elastomers used in bioengineering applications are however typically compressible, some very much so. However, the literature addressing experimental determination of ν for important soft materials such as polyacrylamide gels (PAH, also sometimes referred to as PAA gels in the literature) and polydimethylsiloxane (PDMS) is surprisingly limited when compared to the literature reporting values of *E* and *G*.^14–18^. For these materials, the Poisson’s ratio ν deviates significantly from 0.5 and attains values ranging between 0.25 and 0.49,^13–15, 19^ depending on the molecular weight and the degree of crosslinking of their constituents.^20^

Attempts to quantify or predict the Poisson’s ratio ν of various polymeric materials have used both experimental approaches as well as theoretical models. For example, the study^14^ by Takigawa *et. al.* used a tensile tester to measure the Poisson’s ratio of 27 wt% PAH gels. The gels were kept hydrated and in isothermal conditions using a water bath. Under applied tensile loads, the stretch ratios parallel and perpendicular to the stretched directions were measured, and the value of ν directly estimated. In the same study, the role of strain rate in impacting the Poisson’s ratio was also investigated by varying the strain rate while keeping the PAH formulation constant. Interestingly this study reported that ν did not depend on strain rates within investigated conditions and reported a nearly constant value ν = 0.457. A more recent study by Javanmardi *et. al.*^15^ also using a similar approach, reported that ν values for PAH increased with increasing acrylamide concentrations and were far from the usually assumed value of 0.5. Specifically, the authors reported values of ν = 0.24, 0.30, and 0.32 for 3%, 4%, and 5% acrylamide concentration, respectively. This contrasts, however, with a third study by Boudou *et. al*.^18^ on PAH, in which ν was reported to be acrylamide concentration independent, with ν = 0.485, 0.486, and 0.474 for 5%, 8%, and 10% acrylamide concentration, respectively. These discrepancies in the current literature highlight the importance of systematically characterizing the mechanical response of PAH and the need for a direct measurement of the Poisson’s ratio. This is especially important in applications to cell mechanobiology where soft hydrogels such as PAH are used as substrates and forces exerted on the material by migrating cells are estimated by measuring deformations and transducing them to stress. For example, techniques based on traction force microscopy (TFM) that are often used to study stresses and associated focal adhesion areas in motile cells, parse the data using analytical linear (or nonlinear) constitutive elasticity models.^2, 15^ Using incorrect values of ν in these models will invariably provide incorrect estimates of stresses exerted by the cells.

Elastic properties of the elastomeric polymer polydimethylsiloxane (PDMS) have also been measured recently. Laser-engraved grid patterns were used to obtain ν, *E*, and *G* by optically measuring the grid pattern distortion when the specimen was mechanically stretched.^16^ Alternatively, ν can also be determined by exploiting thermal expansion and measuring surface deformations. Using this approach, Mueller *et al.*^17^ estimated ν, and reported values for the silicone elastomers Sylgard 184 and Sylgard 182 of ν = 0.495 and ν = 0.4974, respectively. These values, however, need to be considered carefully, as the time and temperature used for curing, in addition to the formulation (such as the base to curing agent ratios), are crucial in determining the final structure and therefore mechanical responses of thermally set elastomers, such as PDMS.^21^

The clear need - evident from reviewing the literature - for the characterization of the elastic and viscoelastic properties of substrates used in mechanobiology studies on mammalian cells extends to studies on prokaryotic cells such as bacteria. A recent study found that biofilms of *Serratia marcescens*, *Pseudomonas aeruginosa*, *Proteus mirabilis*, and *Myxococcus xanthus*, have all been found to expand faster on stiffer PAH substrates than on softer ones. TFM measurements showed that the colonies generated transient forces that are correlated over length scales much larger than a single bacterium, and that the magnitude of these forces increases with increasing substrate stiffness.^2^ Understanding these trends requires a clear quantification of the (compressional and shear) stress fields in the underlying soft substrates and relating them to intrinsic substrate elastic properties.

Additional motivation comes from theoretical models and simulations that probe how cells sense soft substrates and interact with each other via substrate mediated elastic communication.^3, 22–27^ Cells act as force dipoles deforming underlying substrates and generating a strain field that can cause nearby cells to reorient and attain energetically favourable configurations. In recent work, we have shown using stochastic agent-based models that the Poisson’s ratio could play an important role in setting the type and range these moderate and short-range biophysical interactions.^28, 29^ Indeed, more recent work suggests that the Poisson’s ratio ν can determine the favorable configurations (both position and orientation) of a pair of dipoles^30, 31^ and direct multicellular network formation on elastic substrates. Using experimentally determined values of the Poisson’s ratio and the elastic moduli rather than approximated values will allow for more realistic theoretical investigations of cell motility, cell-cell interactions, and related mechanobiology problems.

Inspired by the simple macroscopic approach of Pelhalm and Wang^26^ to estimate the Young’s modulus *E* of PAH, we describe herein a similarly simple macroscopic method employing tensile tests to measure directly both *E* and ν. We use our methodology to achieve two objectives. First, we directly characterize the Poisson’s ratio and the two elastic moduli for (three formulations of) PAH, and (three formulations of) PDMS, both of which are relevant substrates for mechanobiology and bioengineering applications. This is done using two approaches. The first approach is using a static tension test from which *E* and ν are directly obtained; the shear modulus *G* can then be calculated assuming the material to be linearly elastic and isotropic. We also use a second independent method based on shear rheology to directly obtain the bulk shear modulus *G* of PAH and PDMS sample. Importantly, we show that the Poisson’s ratio may vary significantly from the value for incompressible materials (ν = 0.5). Our second objective is to compare the values of the shear moduli from the two independent methods and cross-correlate these values with existing literature. Surprisingly, we find a high degree of agreement between shear rheology and macroscopic tension tests for PAH but not PDMS.

Taken together, our study emphasizes the importance of accurate characterizing *E* and ν of gels and elastomers, rather than assuming the incompressible value, especially for use in mechanobiology studies. Our method provides an easy, accessible, and affordable means to achieve this characterization using materials and means commonly found in most laboratories.

## MATERIALS AND METHODS

### PAH rod sample preparation

40 % acrylamide (Sigma Aldrich), 2% bis-acrylamide (Sigma Aldrich), ammonium persulfate (APS) (Invitrogen), and tetramethylethylenediamine (TEMED) (Thermo-Fisher Scientific), and milli-Q water of the indicated volume shown in Table 1 were mixed together in a 15 mL conical tube, starting with the larger volumes.^6, 32, 33^ The mixture was then degassed for 10 minutes. Next, 5 µL of a 10 wt% APS previously prepared and stored at −20°C was added and the entire solution and vortexed. Next, 0.5 µL of TEMED was added and the solution then vortexed again. Lastly, the resulting solution was cast, approximately 7 mL, into disposable straws 6 mm in diameter, with one end sealed with parafilm and held under active vacuum (120 torr) during the curing time of approximately 30 minutes. The PAH rods were removed from the mold and immersed in excess milliQ water (18.2 MΩ cm, TOC < 5 ppm) and allowed to swell overnight at 4 °C, enough to fully swell.^34^

**Table 1.** Formulations used to fabricate PAH. Soft, intermediate, and stiff PAH nomenclatures are based on previously reported protocols.

| **Nomenclature** | **Acrylamide/Bis-acrylamide (%/%)** | **Acrylamide from 40 wt/v% stock (μL)** | **Bis-acrylamide from 2 wt/v% stock (μL)** | **Water (μL)** | **TEMED (μL)** | **APS (μL)** |
| --- | --- | --- | --- | --- | --- | --- |
| Soft | 5/0.3 | 125 | 150 | 725 | 0.5 | 5 |
| Intermediate | 8/0.25 | 200 | 100 | 700 | 0.5 | 5 |
| Stiff | 8/0.48 | 200 | 240 | 560 | 0.5 | 5 |

### PDMS rod sample preparation

Polydimethylsiloxane (PDMS) rods of 50:1, 20:1, and 10:1 (base: curing agent) ratios were created by mixing the appropriate base with curing agent solution using the SYLGARD 184 silicone elastomer kit (see Table 2) and stirred for 5 minutes or so until fully mixed. The entire solution was degassed for 30 minutes under vacuum, ensuring that bubbles in the solution were completely removed. 3.5 mL of degassed and pre-mixed solution was pipetted into disposable straws 6 mm in diameter. One straw end was sealed by using a binder clamp. The filled straw was then immediately placed in an oven at 65 °C for 24 hours for cross-linking. No ramping temperature rate was used.

**Table 2.** Receipts used to fabricate elastic PDMS.

| **Nomenclature** | **Base:Curing agent ratios** | **Base (g)** | **Curing agent (g)** |
| --- | --- | --- | --- |
| 10:1 | 10:1 | 10 | 1 |
| 20:1 | 20:1 | 20 | 1 |
| 50:1 | 50:1 | 50 | 1 |

### Static tensile tests

Fully swollen PAH and PDMS rods were first prepared by hydrating them in water. These were then carefully mounted on the stretcher device, as shown in Figure 1(a), with two clamps attached to each end. The average diameter of our water-swollen PAH and PDMS rods were 6.64 ± 0.65 mm and 6.54 ± 0.07 mm respectively. Two and sometimes three stains, were made (inked) using a permanent marker and used as fiducial markers as seen from Figure 1(b). These were made far from the clamps and close to the center of the rods. Inked gels were then subjected to static tensile forces by applying dead weights (see SI Tables 1 and 2) on the lower end of the PAH rods. Using a digital single-lens reflex camera (Nikon, D750) with a macro lens (Nikon, AF-S Micro Nikkor 105) mounted on a tripod, pictures were taken via a wireless intervalometer to prevent mechanical drift. The PAH and PDMS rods were imaged once with each incremental step of dead weights added, ensuring that the fiducial markers were within the field of view. The camera remained static. Images were subsequently post-processed in FIJI (NIH).

**Figure 1.**
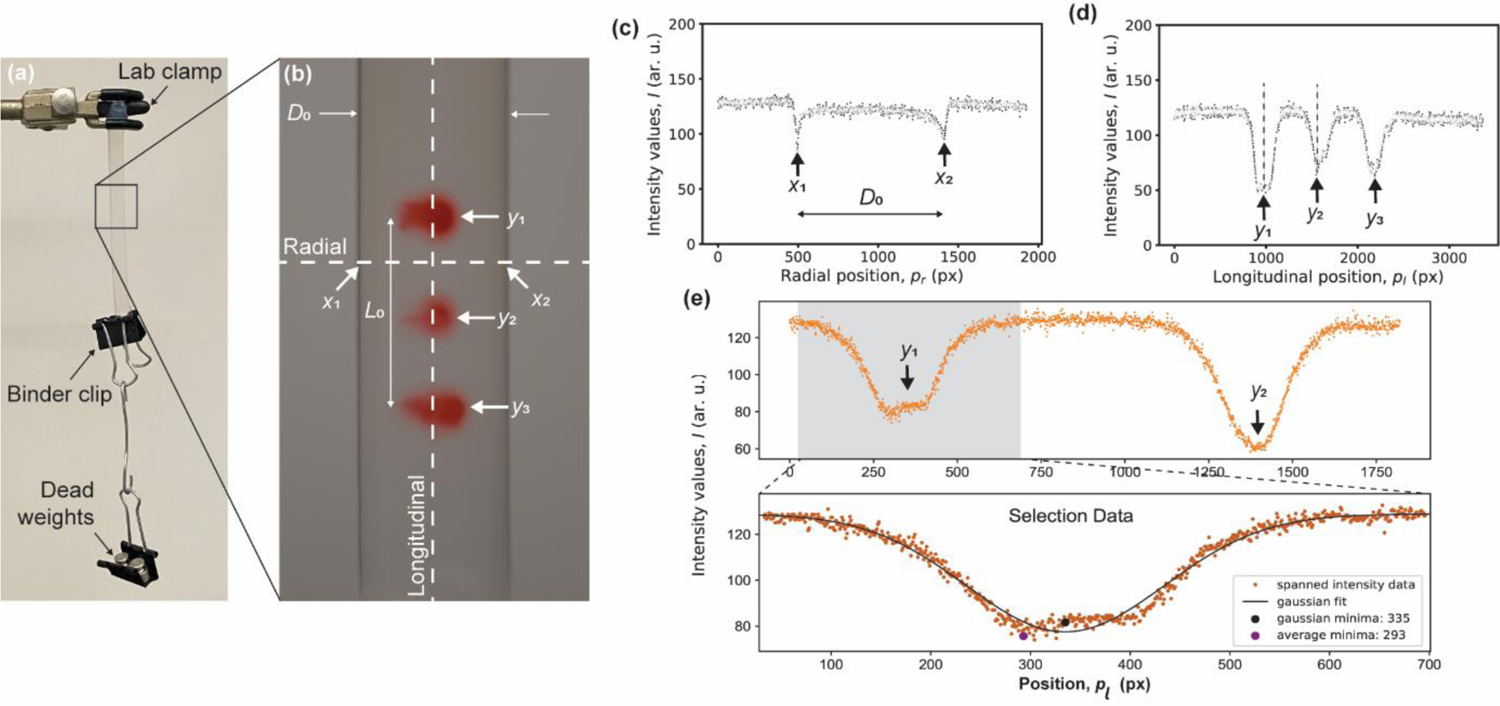
(a) Static tensile test configuration used to extract Poisson’s ratio ν and Young’s modulus *E*. (b) Fiducial markers and line orientations to extract intensity profiles. (c) and (d) Line intensity profiles used to extract diameter (radial) and length (longitudinal) dimensional changes. (e) Intensity Profile Analyzer (IPA) region selection to extract the peak position of the intensity profile.

Intensity profiles – both horizontal (radial) and vertical (longitudinal) – were subsequently traced and analyzed to extract the edge pixel positions via an in-house intensity profile analyzer (IPA) code. Implementation details and use can be found in SI. From these, we obtained the diameter, *D*_i_ (in pixels), and the center-to-center distance of the fiducial markers, *L*_i_ (in pixels), every time a dead weight (indexed by *i*) was added, Figure 1(c-d). The diameter *D*_i_ and distance *L*_i_ were then used to compute the radial and longitudinal engineering strains, ε_R_ and ε_L_defined in equations 1a and 1b:

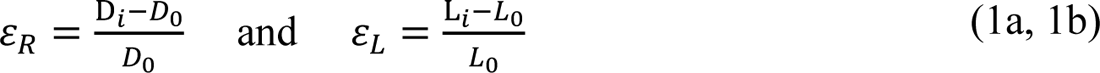

where *D*_0_ and *L*_0_ are the initial diameter and center-to-center fiducial marker length, respectively. The engineering stress σ_i_ was also calculated using equation 2:

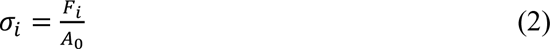

where F_i_ is the tensile force imposed by each dead weight (index *i*) increment and *A*_0_ is the initial cross-section of the rod.^35^

### Poisson’s ratio measurements

To calculate the Poisson’s ratio ν, the radial strain ε_R_ was plotted as a function of the longitudinal strain ε_L_. A linear regression model was then used to calculate the slope, thus extracting the Poisson’s ratio 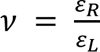, as illustrated in Figure 2(a).

**Figure 2.**
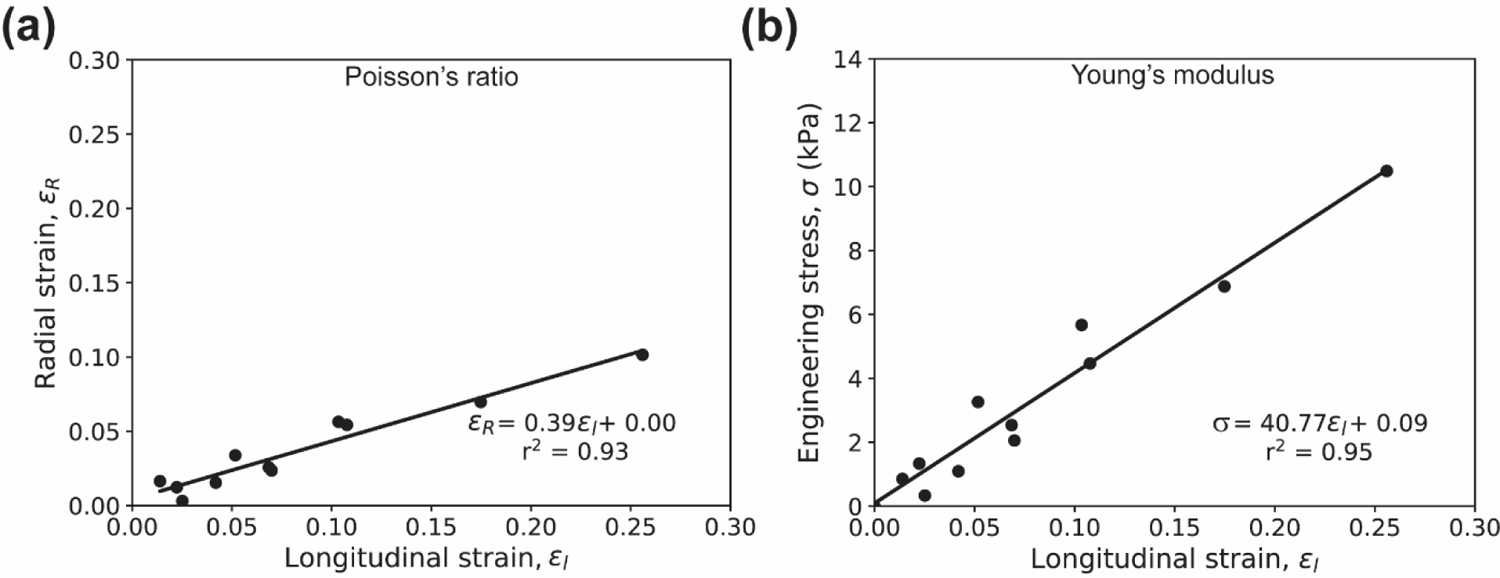
Representative scatter plots and linear regressions for used to extract (a) the Poisson’s ratio and (b) Young’s modulus values.

### Young’s modulus measurements

To calculate the Young’s modulus *E*, the engineering strain σ_i_ was plotted as a function of the longitudinal strain ε_L_. A linear regression model was used to calculate the slope, and the modulus evaluated using 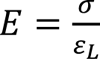, as illustrated in Figure 2(b). Maximum applied axial strains (using weights summarized in SI Table 1) were ≈ 0.3, for the three PAH formulations, which we call soft, intermediate, and stiff PAH, respectively (see Table 1). The maximum applied strains were ≈ 0.9, 0.5, and 0.15 for 50:1, 20:1, and 10:1 for PDMS (using weights summarized in SI Table 2), respectively.

A small pre-stress of 0.3 kPa was applied to PAH samples, 1.2 kPa to 50:1 PDMS samples, and 29 kPa to 20:1 and 10:1 PDMS samples, before starting to quantify strain values. This pre-stress stabilized the rod specimens. Typically, dog-bone shape specimens are recommended for uniaxial tests, to reduce the influence of stress concentrations induced by loading grips at each end of the specimen. Here however, the specimen geometry was kept as simple as possible, and since we applied small strains, the effects of the boundary are negligible.

### Rheology

PAH were created as described by Tse and Engler.^32^ Bulk shear rheological behaviors of PAH and PDMS were characterized using a rheometer (Anton-Paar MCR-302e). The attachment geometry used was a sandblasted stainless-steel parallel plate (PP-25/S), 25 mm in diameter for all measurements. The sandblasted surface was chosen to avoid slippage of the samples in contact with the steel surfaces. The rheology experiments for all gel samples were conducted at a temperature of 25 °C.

For PAH samples, a total of 510 μL of premixed PAH precursor solution (see Table 1) was pipetted onto the bottom plate. Once in place, the sandblasted top parallel plate was slowly lowered until reaching a gap of 1.00 mm, ensuring that the gel sample bridged the top plate, and filled the cavity entirely without voids. PAH samples were then left to polymerize for 30 minutes.^32^ Once polymerization was complete, excess liquid was wiped and the gap was further lowered to 0.990 mm, corresponding to 1% compression strain in the axial direction. We find that the PAHs cast between the parallel plates are very close to their fully swollen state, SI Figure 3 and SI Table 5.

**Figure 3.**
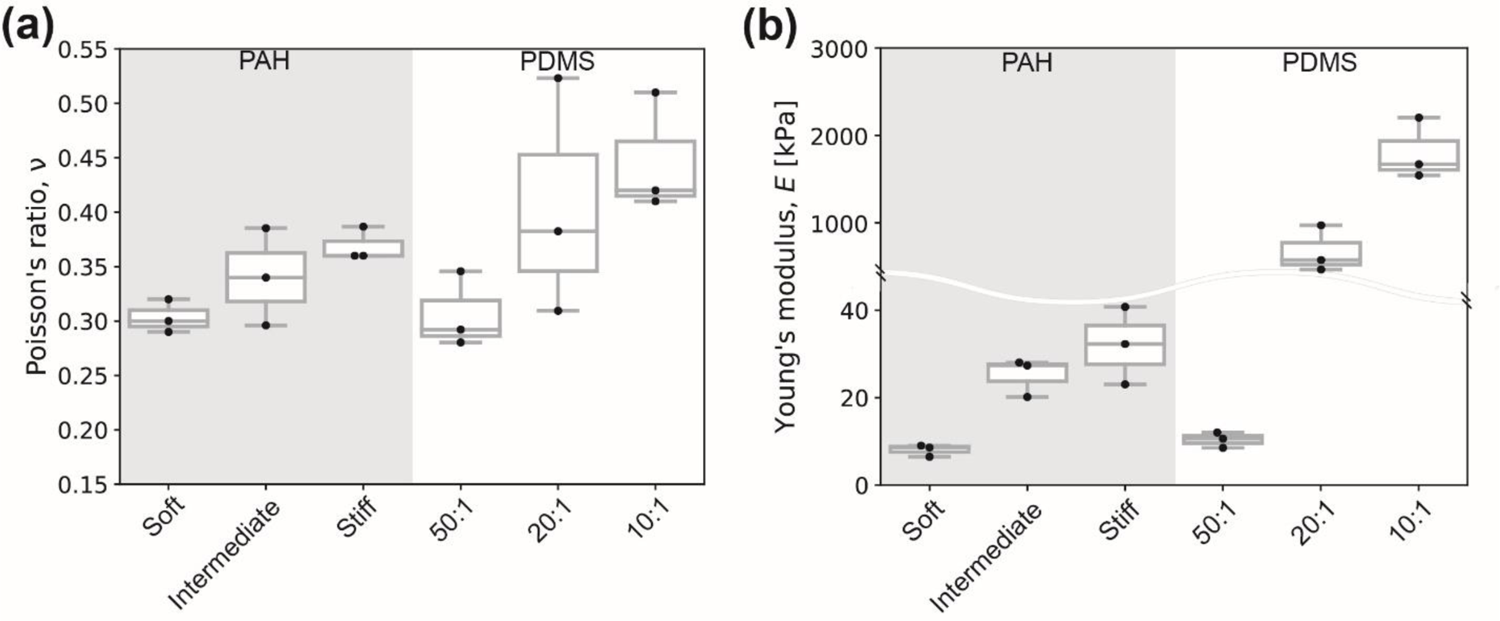
Elastic constants quantified via static tensile tests. (a) Poisson’s ratio values and (b) Young’s moduli of different PAH and PDMS formulations.

For PDMS samples, a total of 2 mL of degassed and premixed base and curing agent solution (see Table 2) was cast onto a 35 mm petri dish lid, resulting in a 1.5 ± 0.3 mm thick film after curing, measured with a vernier caliper. Discs were cut using a 25 mm stainless steel diameter hole punch to match the dimensions of the 25 mm sandblasted rheometer top spindle attachment. Before conducting measurements, the top spindle was lowered while continuously monitoring the normal force measured by the rheometer. We ensured that the normal force was slightly greater than zero (≈ 0.1 N) to ensure the spindle was in contact with the sample. The gap was reduced to achieve a 1% compression strain in the axial direction. This pre-strain combined with the sandblasted surface of the parallel plates avoids sample slippage and ensures contact during the shear tests.

The static shear modulus *G* may be obtained as the limiting value (in the limit of infinitesimally small strain or frequency) of the dynamically measured storage modulus *G*’. The storage moduli *G*’ were measured as a function of angular frequency *ω*, and in an independent set of experiments on the shear strain *γ*. To also obtain an understanding of the range over which the material behaves as a linear elastic system, we probed the samples over a wide range of frequencies and strains. For the frequency sweep tests, the frequency *ω* varied between 0.1-10 rad/s (or equivalently 0.0159-1.59 Hz) at constant shear strain *γ* of 1%. The value of the shear strain (1% shear strain) was chosen so that the material response was linear, and facilitates the comparison of our results with published literature values.^32^ For the shear sweep experiments, the shear strain *γ* was varied between 0.1-10% at constant frequency *ω* of 6.28 rad/s (or equivalently 1 Hz).

### Data analysis and statistics

Data analysis was performed using in-house python scripts (available for download in https://gitlab.com/randresen/facile-determination-of-the-poisson-s-ratio-and-young-s-modulus-of-polyacrylamide-gels-and-polydimethylsiloxane/-/tree/main/. In the plots shown, box and whisker plots indicate the median, with each individual data point corresponding to an independent measurement. Scatter plots show the mean and standard error of the mean.

## Results

### Static tensile Poisson’s ratio and stiffness of PAH and PDMS

The elastic parameters measured from static tension tests for PAH and PDMS are shown in Figure 1 and other related details are summarized in SI Table 3. We observed that the Poisson’s ratio ν of both materials tested increased with increasing initial polymer volume fractions (for PAH) or crosslinking degree (for PAH and for PDMS), indicating that the gels were becoming less compressible, Figure 1(a). For soft, intermediate, and stiff (expected from the fabrication protocol^6^) PAH gels, ν values were 0.30 ± 0.01, 0.34 ± 0.03, and 0.37 ± 0.0.1, respectively. For 50:1, 20:1, and 10:1 PDMS, ν values were 0.31 ± 0.02, 0.41 ± 0.06, and 0.45 ± 0.03, respectively. The Young’s modulus *E* followed a similar qualitative trend as what we just described for the variation in ν of PAH and PDMS. We note that these trends are consistent with classical polymer theory.^36^ As expected, and widely reported in literature,^6, 7, 15^ the Young’s modulus *E* of PAH increased with increasing acrylamide concentration (polymer volume fraction), with *E* values ranging from 8.0 ± 0.8 kPa, to 25.2 ± 2.5 kPa, and 32.0 ± 5.1 kPa for soft, intermediate, and stiff PAH, respectively. The results are in good agreement with *E* values obtained via AFM nanoindentation^6^ and other macroscopic studies.^26^

Increasing the curing agent content of PDMS results in a higher degree of cross-linking in the polymeric matrix, and therefore increasing *E*. The static tensile test *E* values measured for PDMS were 10.4 ± 0.75 kPa, 667.4 ± 153.8 kPa, and 1802.0 ± 202.3 kPa for 50:1, 20:1, and 10:1 mixing ratios, respectively. These results are also in excellent agreement with other studies reporting on tensile characterization of PDMS using more sophisticated approaches, such as a universal testing machine or dynamic mechanical analysis.^9, 37–39^

To ensure elasticity of PAH and PDMS, we quantified the Young’s modulus during unloading, that is, by removing dead weights, and obtained very similar values for all conditions tested, SI Figure 1(a) and (b) and SI Table 4.

### Rheology of PAH and PDMS

To characterize the shear modulus *G*’ of the soft, stiff, and intermediate PAH gels, and of the 50:1, 20:1, and 10:1 PDMS gels, we conducted shear rheology measurements. Note that the zero-frequency shear modulus *G* may be obtained by examining the frequency dependence of the rheologically measured *G*’. As previously stated, we report *G* as the limit of *G*’ as *ω* or γ tends to zero.

We first present results for the PAH gel samples probed by rheology. Figure 4(a) shows the log-log curves of *G*’ of soft, intermediate, and stiff PAH as a function of shear strain, *γ*. Our results confirm that for small to moderately small shear strains (in the range 0.01-10%), all PAH samples behaved as a linear elastic solid, as suggested by the near-constant (with negligible linear slope) values of *G*’. The initial average *G*’ values measured from the shear strain sweeps were 2.1 ± 0.2 kPa, 7.2 ± 0.8 kPa, and 10.7 ± 1.0 kPa, for soft, intermediate, and stiff PAH respectively. Figure 4(b) shows the frequency dependence of *G*’. Our results reveal that for small to moderately small angular frequencies (in the range 0.1-10 rads/s), PAH behaves as a linear elastic solid. The first measured *G*’ value for soft, intermediate, and stiff PAH measured from the frequency sweep tests were 2.7 ± 0.1 kPa, 6.6 ± 0.4 kPa, and 10.2 ± 1.1 kPa, respectively.

**Figure 4.**
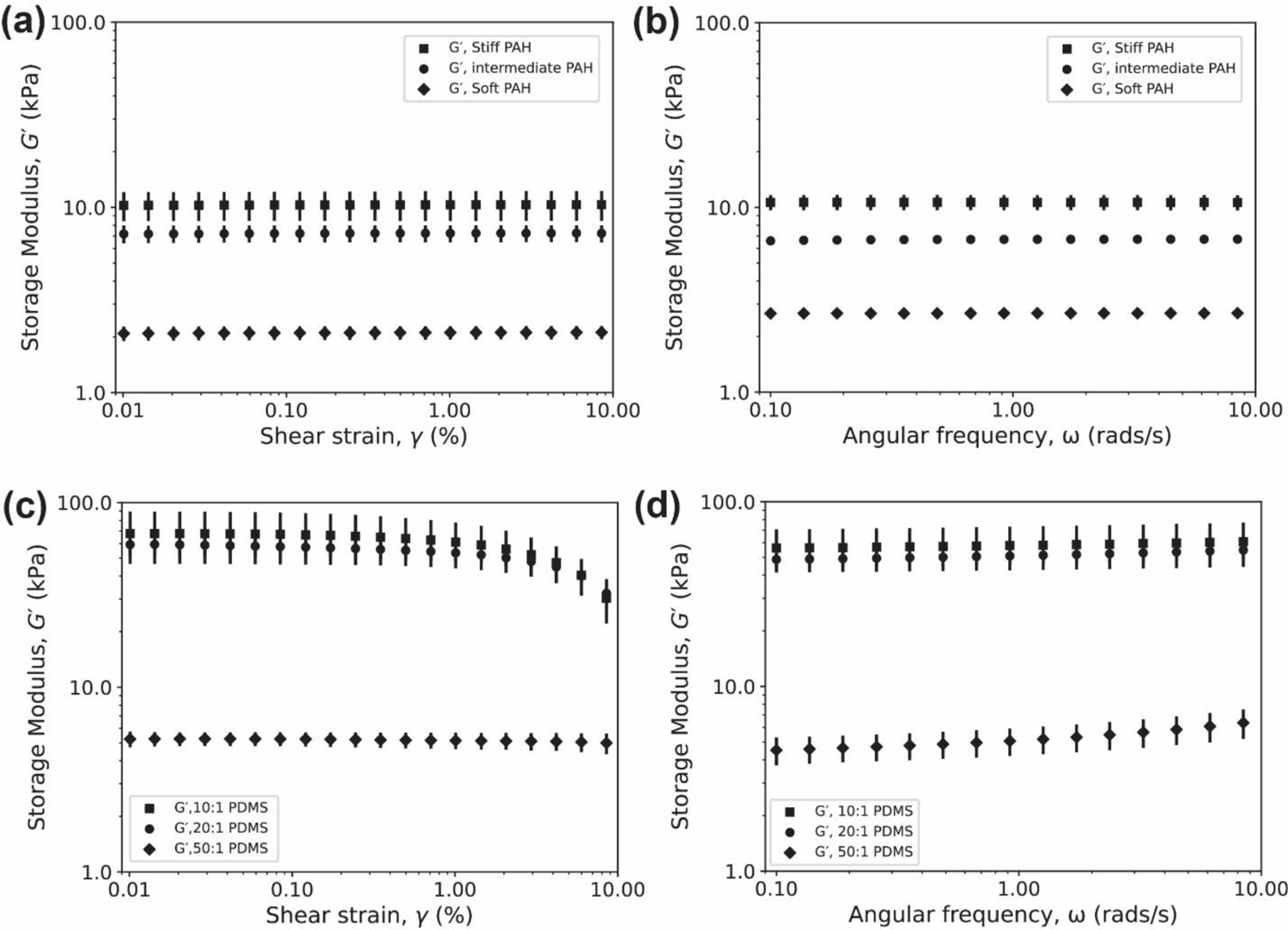
(A and C) Shear strain dependency of the dynamic storage modulus (*G*’) at a constant angular frequency of 6.28 rads/s (or 1 Hz). (B and D) Frequency dependency of the storage modulus (*G*’) at a constant shear strain of 1%. Three independent and new samples per condition are reported. Shown is the mean and the standard error of the mean.

We further quantified the effect of compressive strain on *G*’ of stiff PAH samples. As seen in SI Figure 2(a)-(b) and summarized in SI Table 5, variations in *G*’ are within experimental error, and did minimally vary with increasing compressive strain.

We next present the results of our studies on PDMS samples. Figure 4(c) shows the log-log curves of *G*’ as a function of shear strain, *γ*. For 20:1 and 10:1 PDMS samples, the linear elastic region indicates that the elastomer has predominantly linear elastic behavior up to a shear strain value of approximately 1%. Above 1% shear strain, 20:1 and 10:1 PDMS samples begin to soften. The *G*’ values as a function of *γ* are 59.2 ± 4.6 kPa and 67.9 kPa, respectively. However, 50:1 PDMS behaved linearly over the full range of shear strains tested (up to 10%), with an average *G*’ value of 5.2 ± 0.3 kPa.

As opposed to PAH, all three PDMS formulations showed different rheological responses between the sweep modes. Figure 4(d) shows the log-log curves of *G*’ as a function of frequency, *ω*. The 50:1 formulation shows stiffening. From an initial value of 4.5 ± 0.5 kPa, *G*’ increased to a value 6.4 ± 1.2 kPa, approximately 40% stiffer at the highest frequency tested. The 20:1 and 50:1 PDMS samples behaved differently. For these, *G*’ remained constant over the range of frequencies tested (up to 10 rad/s), with *G*’ values of 48.7 ± 2.8 kPa and 56.1 ± 8.5 kPa, respectively. Overall, the PDSM samples deviated from linearity depending on the shear test (*i.e.*, strain sweep or frequency sweep), suggesting the importance of *a priori* knowledge of the application intended for PDMS.

We further quantified the effect of compressive strain on *G*’. As seen in SI Figure 2(b)-(c), *G*’ increases with increasing compressive strain. This result emphasizes the importance of properly reporting characterization parameters, as the quantified values will vary significantly.

### Impact of assumed and measured Poisson’s ratio on shear stiffnesses

As mentioned previously, our first goal was to directly measure the Poisson’s ratio of the PAH and of the PDMS formulations. Our second goal was to cross-evaluate and compare *E* obtained from the static tension test, with that estimated from shear rheology by using *G*. We convert from the Young’s modulus to the shear modulus by assuming the continuum relationship valid for a linear isotropic elastic material (equation 3) and using the appropriate value of the Poisson’s ratio,

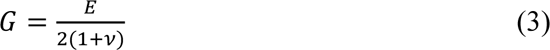

We use equation 3 to calculate the shear modulus by i) assuming incompressibility, and ii) by using the actual (measured) value of the Poisson’s ratio. This allows us to quantify variations in the predicted value of *G*, resulting from the approximation. We perform this analysis first for PAH, followed by PDMS.

### Comparison between rheology and static tensile tests of PAH

Most previous studies on PAH as substrates, assume that it is incompressible with a Poisson’s value of 0.5. We consider the first measurable *G*’ value (*i.e.*, at the lowest frequency and/or the lowest shear-strain) to be equivalent to the shear modulus *G*. Figure 5(a)-(b) shows the cross-evaluation between *G*_Tension_ (deduced from *E* measured from static tensile tests) and *G*_Rheology_ (deduced from *G*’ measured from shear strain sweep tests), and Figure 5(c)-(d) the cross-evaluation between *G*_Tension_ and *G*_Rheology_ (this time deduced from frequency sweep tests). The dashed lines in Figures 5(a) and (c) indicate a perfect cross-correlation between the two approaches used to characterize the elastic responses of PAH, with a slope of 1. Assuming that the PAHs are incompressible, ν = 0.5, the cross-correlation between *G*_Tension_ and *G*_Rheology_ for *G*’ values determined from the shear strain sweeps, Figure 5(a), deviates to a lower slope value of 0.95, suggesting that the PAHs are slightly softer when characterized via static tension tests when compared against shear rheology. If the experimentally determined ν is used for the cross-correlation between *G*_Tension_ and *G*_Rheology_, we find that the slope takes a value of 1.02, indicating that the PAHs are slightly stiffer than when characterized via static tension tests when compared against shear rheology. The differences, however, for both ν, ideal incompressible and measured are relatively small and within experimental error. For the case of *G*_Rheology_ (equivalently *G*’) values obtained from the frequency sweep tests, for both ν, ideal incompressible and measured, the cross-correlation between *G*_Tension_ and *G*_Rheology_ deviates to a larger slope value of 1.06 and 1.14, respectively, indicating that the PAHs are slightly stiffer when characterized via static tension tests when compared against shear rheology.

**Figure 5.**
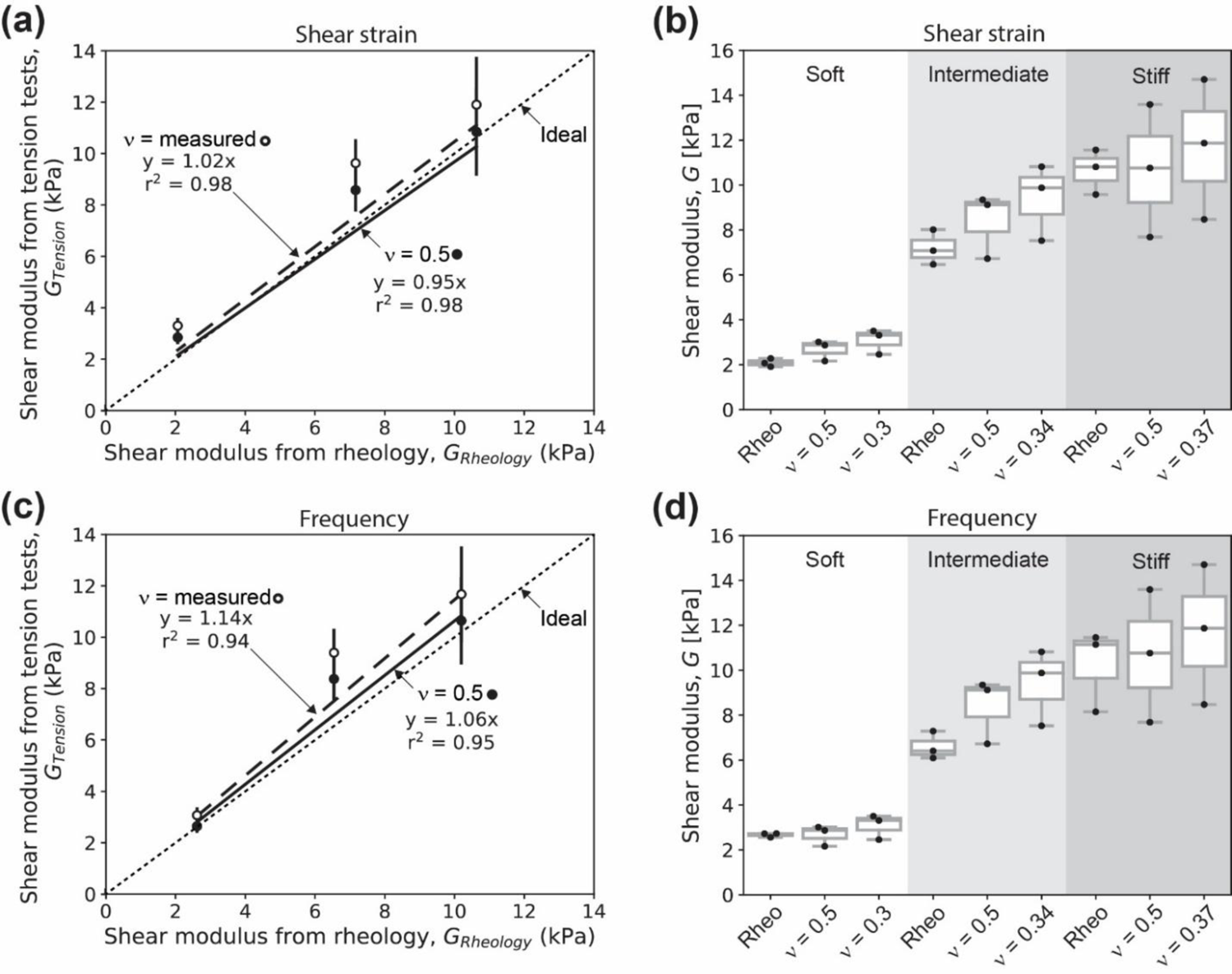
Cross-correlation of Young’s modulus (*E*) and shear modulus (*G*) as a function of Poisson’s ratio (ν). The dotted lines represent a 1:1 ideal correlation. (a) and (b) Cross-correlation comparison for shear strain sweeps at constant *ω* of 1 rad/s. (c) and (d) Cross-correlation comparison for frequency sweeps at constant 1% compressive strain of PAH samples. Error bars represent the standard error of the mean.

Overall, we observe small deviations (within experimental error) in the cross-correlation agreement between *G*_Tension_ and *G*_Rheology_ estimated using both an idealized incompressible value of ν and exactly measured value. This, however, does not imply that it is a valid assumption to always consider ν as incompressible, as further discussed in the Discussion section and elsewhere.^13, 15^ This is especially important when we consider dynamic effects, specific to mechanobiology studies on cell motility on biomimetic hydrogel surfaces. For instance, the deformation fields induced in the substrate due to the contractile and time-dependent stresses in focal adhesion regions depend crucially on the Poisson’s ratio.^28, 29, 40^ A second instance where Poisson’s ratio may play an important role is in the rate-dependent deformation of initially planar hydrogels of finite stiffness. Here, axial and normal deformations are coupled via the Poisson’s ratio, and together determine the time dependent forces felt by the indenter. Such rate-dependent deformations are not static, and involve significant poroelastic effects.^41^

### Comparison between rheology and static tensile tests of PDMS

Next, we extended the above-described approach to analyze the cross-correlation of elastic constants obtained between static tensile tests and shear rheology for PDMS rod. We note that unlike PAH, PDMS is an elastomer. We asked whether the cross-correlation of the two bulk characterization modes (tension *vs*. compression with shear) was in good agreement, as seen for PAH. Figure 6(a)-(b) shows the cross-evaluation between *G*_Tension_ (deduced from *E* measured from static tensile tests) and *G*_Rheology_ (deduced from *G*’ measured from shear strain sweep tests), and Figure 6(c)-(d) the cross-evaluation between *G*_Tension_ and *G*_Rheology_ (this time deduced from frequency sweep tests). The dashed lines in Figures 6(a) and (c) indicate a perfect cross-correlation between the two approaches used to characterize the elastic responses of PDMS, with a slope of 1. Assuming that the PDMS samples are incompressible, ν = 0.5, the cross-correlation between *G*_Tension_ and *G*_Rheology_ for *G*’ values determined from the shear strain sweeps, Figure 6(a), deviates considerably to a larger slope value of 7.60, suggesting that the static tension tests on the PDMS results in a stiffer response and a larger value for the modulus than that estimated via shear rheology. If the experimentally determined ν is used for the cross-correlation between *G*_Tension_ and *G*_Rheology_, we find that the slope increases even further, to a value of 7.92, indicating that the PDMS samples seem even stiffer. The differences for both ν, ideal incompressible and measured are relatively small and within experimental error as well.

**Figure 6.**
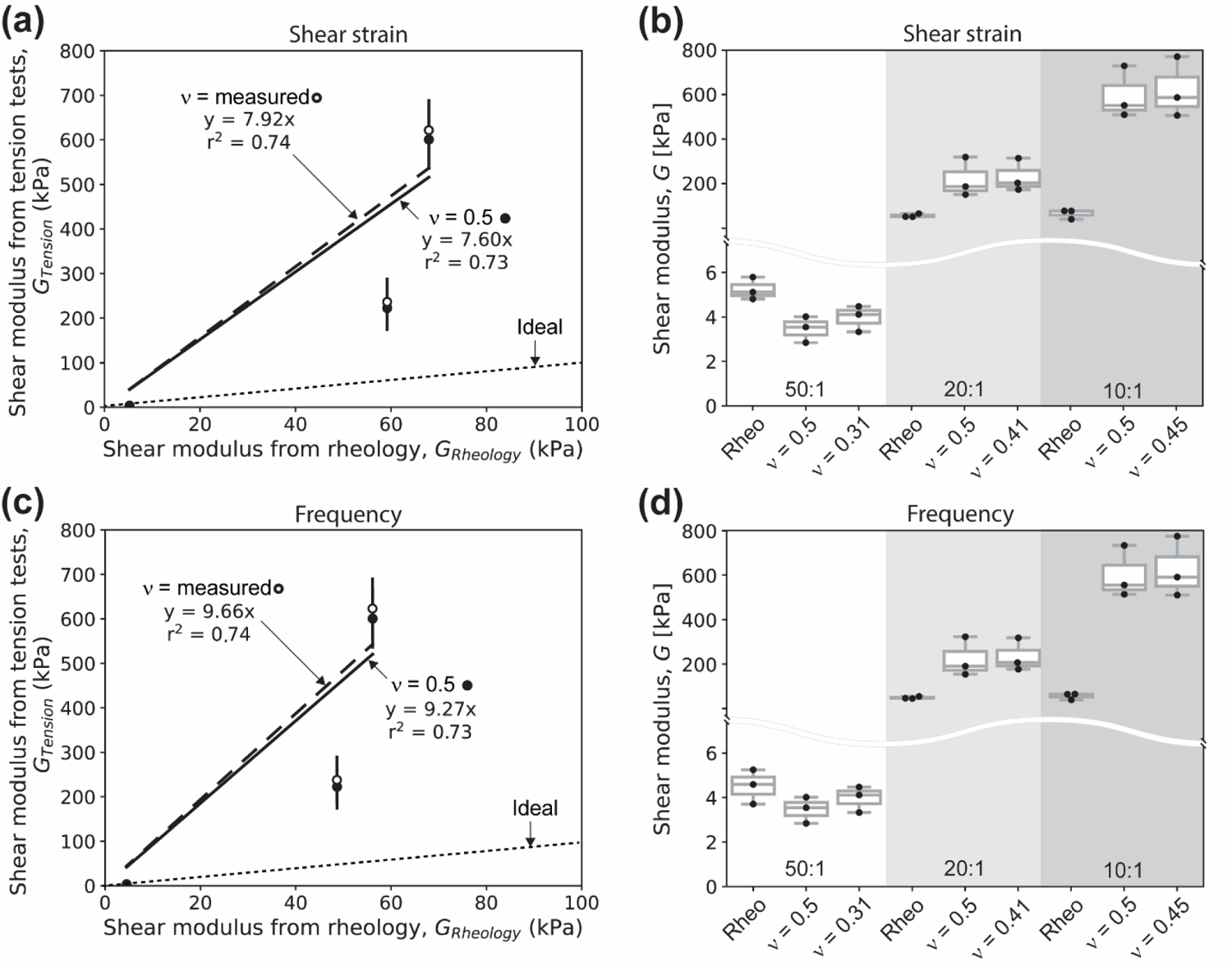
Cross-correlation of Young’s modulus (*E*) and shear modulus (*G*) as a function of Poisson’s ratio (ν). The dotted lines represent a 1:1 ideal correlation. (a) and (b) Cross-correlation comparison for shear strain sweeps at constant *ω* of 1 rad/s. (c) and (d) Cross-correlation comparison for frequency sweeps at constant 1% compressive strain of PDMS samples. Error bars represent the standard error of the mean. Note that in panels (b) and (d), the y-axes have a range break.

## Discussion

The primary goal of this work was to evaluate the Poisson’s ratio of gels and polymers such as PAH and PDMS relevant to a range of disciplines, including mechanobiology, bioengineering, biomaterials, among others. Many experimental studies that interpret data obtained using these gels and elastomers as substrates, as well as other computational studies assume that the PAH is incompressible and thus ν = 0.5. Here, we corroborate that this assumption is not correct, and that ν of PAHs increases with increasing polymer volume fraction and crosslinking degree, as previously reported^15^ and as predicted by scaling concepts of polymer physics.^20, 42^ The static tensile test describe herein obtains bulk properties. However, these can be scale dependent. For instance, approaches such as embedded microneedles actuated with external magnetic fields yield different values, emphasizing the importance of the scale of the characterization.^43^ Our ν values for PAH and others^14, 15, 43^ are summarized in Figure 7(a). We observe a similar behavior for PDMS, that is, ν increased with increasing degree of crosslinks. Our ν values for PDMS and others^16, 17, 44^ are summarized in Figure 7(b).

**Figure 7.**
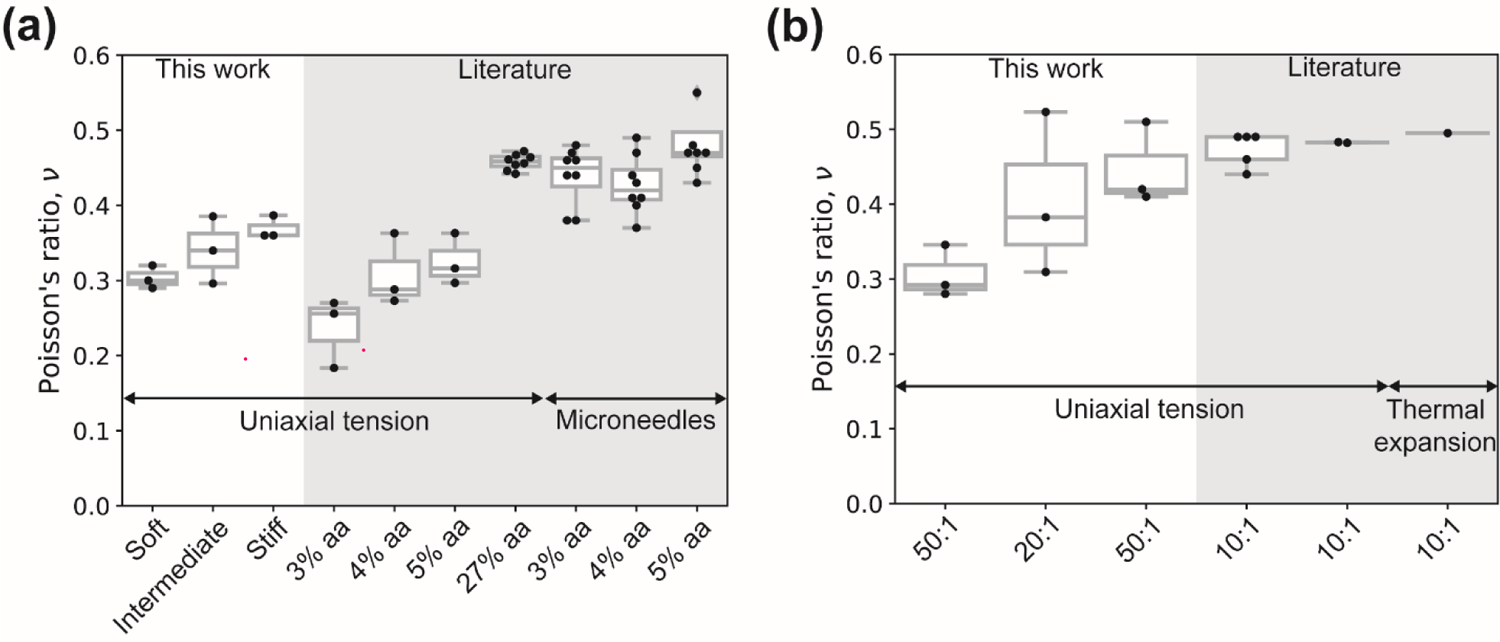
Comparison of Poisson’s ratio ν values from this work and other works for (a) PAH and (b) PDMS. All PDMS values reported correspond to Sylgard 184.

Quantifying these two elastic constants is of crucial importance for experimental platforms, such as traction force microscopy (TFM), the most widely employed contractile force measurement approach for adherent cells, which relies on knowledge of the force-displacement relationship and mechanical properties of the substrate.^1, 2, 45–49^ In a recent study, Javanmardi *et al.* demonstrated that applying the correct ν is crucial for accurate force reconstruction employing TFM approaches.^15^ One advantage of our approach is that it is extremely simple. Specifically, a single experiment can be used to extract values of both, Young’s modulus *E* and Poisson’s ratio ν, for soft materials relevant to mechanobiology.

In general hydrogel networks due to their various cross-linking properties, and the fluid (water or a biological fluid) permeating them, have both viscoelastic and viscoplastic properties. Two main types of the crosslinks used in formulating the hydrogels are chemical crosslinks and physical crosslinks which both confer structure and elastic properties but contribute differently in terms of both frequency dependent and static properties. Chemical crosslinks, for instance in PAH due to irreversible chemical bonding provide greater rigidity and enable static pre-stressed states. Physical crosslinks are comparatively weak and may be temporary and allow for relaxation of imposed stresses. During the deformation of PAH, hydrogen bonds may form and break depending on how close different polymeric chains get. Additionally, other temporary physical crosslinks such as chain entanglements confer some amount of strength to the hydrogel; they may however slip and unentangle under stress. Transient cross-links may significantly contribute to dissipation (and thus to the viscous loss modulus) when hydrogels are subject to frequency dependent deformations. Finally, hydrogels are permeated by a fluid. In the nearly (fully) swollen state, the osmotic pressure due to the fluid content supports the polymer network/scaffold and provides a means to support and balance externally imposed stresses.

Usually, the analysis and interpretation of the elastic response of materials such as metals is assumed to be the same in response to small tension and compression strains.^50^ This is not the case for soft polymeric networks, in which tension-compression asymmetry (TCA) is commonly found.^51^ For instance, Young’s modulus of hydrogels and elastomers evaluated from tension measurements can differ from compression measurements by orders of magnitude.^51^ While both PAH and PDMS have chains bridged by chemical and physical bonds, particularly at low crosslinking densities, PAH (an hydrogel) consists of loosely crosslinked networks permeated by water molecules, and PDMS (an elastomer) consists of chains that are more severely restricted due to increased crosslinking. This difference has a tremendous impact on molecular mechanisms affecting network responses (to stress) such as viscoplastic flow of junctions^52^ or the formation of transient reversible physical bonds such as hydrogen bonds. Furthermore, when fully swollen hydrogels are subject to deformations (that impact local volume conservation and pore deformations) such as compression or tension, water is forced to flow through the pores; the permeability of the network enabling this flow is controlled by the size of the mesh, denoted by *ξ*. If *ξ* becomes small, as is the case for our PAH, SI Table 7, the osmotic pressure, and hence the shear modulus (or equivalently, the Young’s modulus) increases. Classical theories due to Flory^53^ and de Gennes^20^ relate shear modulus *G* (related to polymer concentration), and osmotic pressure Π, via the relationship *G ≈ Π = k_B_^T^/ξ*^3, 20^ where k_B_is Boltzman’s constant, *T* the absolute temperature, and *ξ* the mesh size. In summary, the mechanical response, be it compression, shear, or tension of PAH will be strongly coupled to *ξ*.^42^ We note that the osmotic pressure may also be written in terms of the concentration of polymer (measured in terms of the Kuhn length for instance), and correlated with the pore size.

Under tension or compression, and for small perturbations (*i.e.*, small strains or small frequencies), fluid is forced through the porous, elastic networks characterized by the mesh size *ξ*. We note from our experiments that static tension tests and shear rheology measurements for PAH show good agreement. Richbourg *et al*. cross-correlated five independent stiffness measurement methods for various polyvinyl alcohol (PVA) hydrogels, also finding very good agreement.^54^ This suggests that for relatively simple hydrogels, that is, hydrogels composed of monomers with minimal side group bulkiness, such as PAH (*i.e.*, acetamide) or PVA (*i.e.*, hydroxy), osmotic pressure combined with viscoplastic flow of junctions could be seen as the main mechanisms governing the mechanical response. Contrarily, PDMS, particularly at 10:1 or 20:1 base to curing ratios, where the network is heavily crosslinked and molecular linkages restricted, little to no viscoplastic flow of junctions takes place, and junctions/crosslinks remain “active” during tension contributing to most of the extensional stress. However, the situation changes under compression when many junctions weaken, and the buckling of polymeric chains causes local softening. In other words, weakening of the junctions causes the polymeric chains to switch to a dangling state, implying that the stresses in the involved chains are reduced significantly. While this hypothesis remains to be tested more carefully, the mechanism described, schematized in Figure 8, explains the experimental observations for PDMS, in which 10:1 samples (with highest crosslinking degree) show a high level of discrepancy between rheology (compression plus shear) and static tension tests. The discrepancy decreases for 20:1, and even more so for 50:1 PDMS samples, consistent with a decrease in the cross-linking degree, and resembling more the hydrogel mode (without the osmotic pressure contribution).

**Figure 8.**
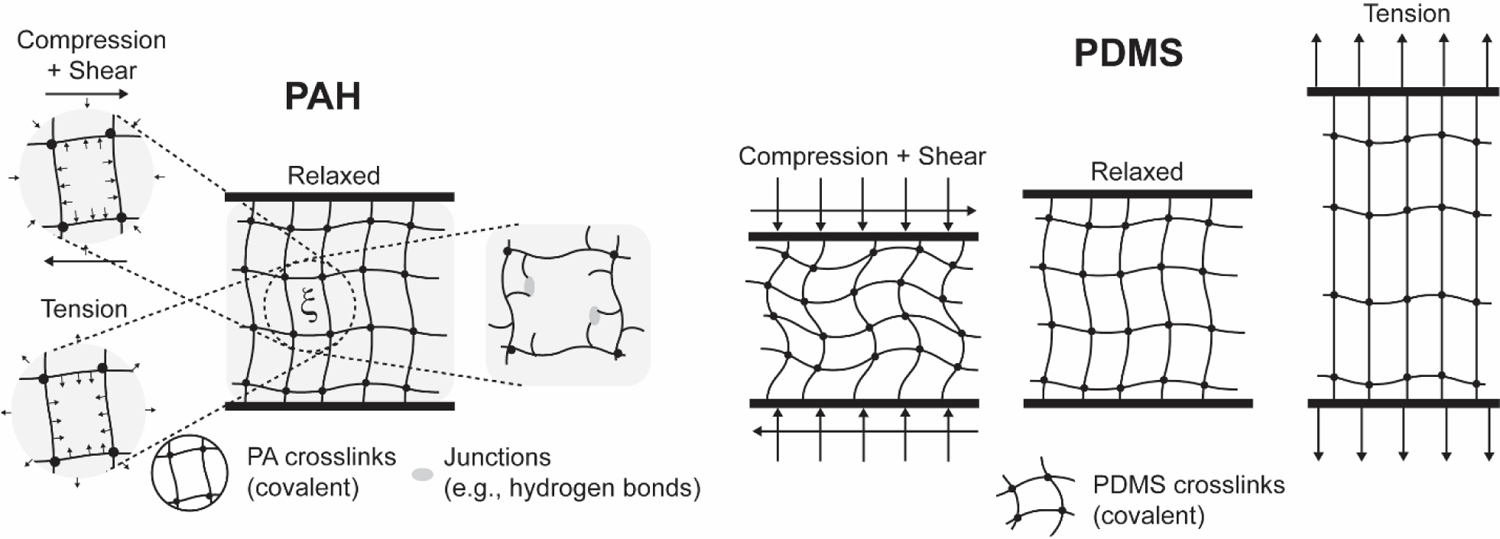
Schematic figures illustrating how osmotic pressure in PAH hydrogels, mesh size, and the presence of viscoplastic junctions (physical crosslinks) mediate mechanical response (left). Figures on the right show the change in network state (with links dangling/buckled under compression, or fully stretched under tension) for a densely crosslinked (10:1, 20:1) PDMS.

## Conclusion

In this study, we characterized the bulk mechanical responses of PAH and PDMS by varying the network compositions. Specifically, we quantified the Poisson’s ratio νand Young’s modulus *E* via static tension tests. We show that the Poisson’s ratio varies from the value for incompressible materials (ν = 0.5), and that its value depends on the crosslinking degree (or alternatively, mesh size). Furthermore, we performed shear rheology to obtain the shear modulus *G* of PAH and PDMS and find that for PAH, the cross-correlation of the elastic constants obtained from the independent methods is in very good agreement, but not for PDMS. Together, our study emphasizes the importance of accurate characterizing *E* and ν of gels and elastomers, rather than assuming the incompressible value, especially for use in mechanobiology studies. Our method provides an easy, accessible, and affordable means to achieve this characterization using materials and means commonly found in most laboratories.

## Supporting information

https://gitlab.com/randresen/facile-determination-of-the-poisson-s-ratio-and-young-s-modulus-of-polyacrylamide-gels-and-polydimethylsiloxane/-/tree/ma

## ASSOCIATED CONTENT

### Supporting Information

The following files are available free of charge: Supporting Information (PDF) Intensity Profile Analyzer (Python scripts)

## AUTHOR INFORMATION

### Author Contributions

The manuscript was written through contributions of all authors. All authors have given approval to the final version of the manuscript. ‡These authors contributed equally.

### Notes

#### ORCID

Ariell M. Smith: 0000-0002-4788-4611

Dominique Gabri Inocencio: 0000-0002-7488-4086 Brandon M. Pardi: 0000-0001-6483-9858

Arvind Gopinath: 0000-0002-7824-3163

Roberto C. Andresen Eguiluz: 0000-0002-5209-4112

## ACKNOWLEDGMENT

A.M.S., A.G. and R.C.A.E. acknowledge funding from the NSF-CREST: Center for Cellular and Biomolecular Machines through the support of the National Science Foundation (NSF) Grant No. NSF-HRD-1547848. A.M.S and R.C.A.E. acknowledge funding from the Tobacco-Related Disease Research Program through the support of University of California Office of the President Grant No. T31KT1583 awarded to R.C.A.E.. A.M.S. and A.G. acknowledge funding from the CAREER grant NSF Grant No. CBET 2047210 awarded to A.G.. D.G.I. acknowledges the fellowship provided by U-RISE through the support of the National Institutes of Health (NIH) Grant No. NIH 1T32GM145511-01.

## ABBREVIATIONS

PAH: polyacrylamide hydrogel

PDMS: polydimethylsiloxane

AFM: Atomic Force Microscopy

TFM: Traction Force Microscopy

TCA: tension-compression asymmetry

PVA: polyvinyl alcohol

