## Supplementary material for "Facile determination of the Poisson’s ratio and Young’s modulus of polyacrylamide gels and polydimethylsiloxane": https://gitlab.com/randresen/facile-determination-of-the-poisson-s-ratio-and-young-s-modulus-of-polyacrylamide-gels-and-polydimethylsiloxane/-/tree/ma

### MATERIALS AND METHODS

#### Static tensile test dead weights

For static tension tests, a series of dead weights from a calibration weights set were added (loading) or removed (unloading) to the PAH rods, SI Table 1, or PDMS rods, SI Table 2. The table summarizes the total dead weight at each weight increment.

**SI Table 1.** Dead weights used for static tension tests of PAH.

| **Stiff PAH** | **Intermediate PAH** | **Soft PAH** | **Weight, *W* (g)** |
|---|---|---|---|
| Pre-stress | Pre-stress | Pre-stress | 1.22 |
| Measurement 1 | Measurement 1 | Measurement 1 | 0.69 |
| Measurement 2 | Measurement 2 | Measurement 2 | 1.38 |
| Measurement 3 | Measurement 3 | Measurement 3 | 3.53 |
| Measurement 4 | | | 5.53 |
| Measurement 5 | Measurement 4 | Measurement 4 | 8.53 |
| Measurement 6 | Measurement 5 | Measurement 5 | 13.53 |
| Measurement 7 | Measurement 6 | Measurement 6 | 18.53 |
| Measurement 8 | Measurement 7 | Measurement 7 | 23.53 |
| Measurement 9 | Measurement 8 | Measurement 8 | 28.53 |

**SI Table 2.** Dead weights used for static tension tests of PDMS.

| **10:1 PDMS** | **20:1 PDMS** | **Weight, *W* (g)** | | **50:1 PDMS** | **Weight, *W* (g)** |
|---|---|---|---|---|---|
| Pre-stress | Pre-stress | 102.6 | | Pre-stress | 4.6 |
| Measurement 1 | Measurement 1 | 200 | | Measurement 1 | 5 |
| Measurement 2 | Measurement 2 | 500 | | Measurement 2 | 10 |
| Measurement 3 | Measurement 3 | 700 | | Measurement 3 | 15 |
| Measurement 4 | Measurement 4 | 900 | | Measurement 4 | 20 |
| | | | | Measurement 5 | 25 |

#### Swelling ratio

Soft, intermediate, and stiff PAH samples were allowed to polymerize in 6 mm diameter disposable straws for 30 minutes as described in the PAH rod sample preparation section. Each sample was cut into multiple sections 12.7 mm in length. Samples were taken from the top, middle, and bottom sections of the rod. To measure the weight change due to water intake, each sample was weighted directly after polymerization (*i.e.*, as cast) and placed in miliQ water. Each sample was weighed at 12 and 24 hrs. We call the mass change the swelling ratio ($SR_{PAH}$), which was calculated using eq. SI 1 for the two time points.

$$SR_{PAH} = \frac{(W_{swollen} - W_{as\ cast})}{W_{as\ cast}} * 100 \qquad \text{(SI 1)}$$

Where $SR_{PAH}$ is the swelling ratio of the PAH sample, $W_{swollen}$ is the weight of the PAH at either 12 hrs or 24 hrs incubation in milliQ water, and $W_{as\ cast}$ is the weight of the PAH after fully polymerized.

**Strain quantification and Intensity Profile Analyzer software (IPA)**

To quantify the PAH and PDSM dimensional changes in pixels, a digital single-lens reflex camera (Nikon, D750) with a macro lens (Nikon, AF-S Micro Nikkor 105) mounted on a tripod to take pictures via a wireless intervalometer to prevent mechanical drift.  The PAH or PDMS rods were imaged once with each incremental step of dead weights added, ensuring that the fiducial markers were within the field of view. The camera remained static. Images were subsequently post-processed in FIJI (NIH). An horizontal line for diameter quantification or a vertical line passing through the center of the fiducial markers were traced, followed by the Plot Profile function under the Analyze menu of FIJI. The pixel and intensity values were saved as *.xls file and further analyzed using the using an in-house Python based tool. The tool, named call Intensity Profile Analyzer (IPA) is available for download at:

https://gitlab.com/randresen/facile-determination-of-the-poisson-s-ratio-and-young-s-modulus-of-polyacrylamide-gels-and-polydimethylsiloxane/-/tree/main/

The "Readme", describing the execution sequence and content is available as a separate file together with the main source code, as well as a sample spread sheet file.

### RESULTS

#### Elastic constants

A summary of the elastic constants for PAH and PDMS samples obtained via static tension tests and rheometry is presented in SI Table 3.

**SI Table 3.** Elastic constants obtained from the static tension tests and shear rheology measurements with 1 % compression strain.

| **Sample** | **Poisson's ratio, *v*** | **Young's modulus, $E_{Tension}$ (kPa)** | **Shear modulus, $G'_{StrainSweep}$ (kPa)** | **Shear modulus, $G'_{FrequencySweep}$ (kPa)** |
|---|---|---|---|---|
| Soft PAH | 0.30 ± 0.01 | 8.0 ± 0.8 | 2.1 ± 0.2 | 2.7 ± 0.1 |
| Intermediate PAH | 0.34 ± 0.03 | 25.2 ± 2.5 | 7.2 ± 0.8 | 6.6 ± 0.4 |
| Stiff PAH | 0.37 ± 0.01 | 32.0 ± 5.1 | 10.7 ± 1.0 | 10.2 ± 1.1 |
| 50:1 PDMS | 0.31 ± 0.02 | 10.4 ± 0.75 | 5.2 ± 0.3 | 4.5 ± 0.5 |
| 20:1 PDMS | 0.41 ± 0.06 | 667.4 ± 153.8 | 59.2 ± 4.6 | 48.7 ± 2.8 |
| 10:1 PDMS | 0.45 ± 0.03 | 1802.0 ± 202.3 | 67.9 ± 12.4 | 56.1 ± 8.5 |

**Validation of elasticity**

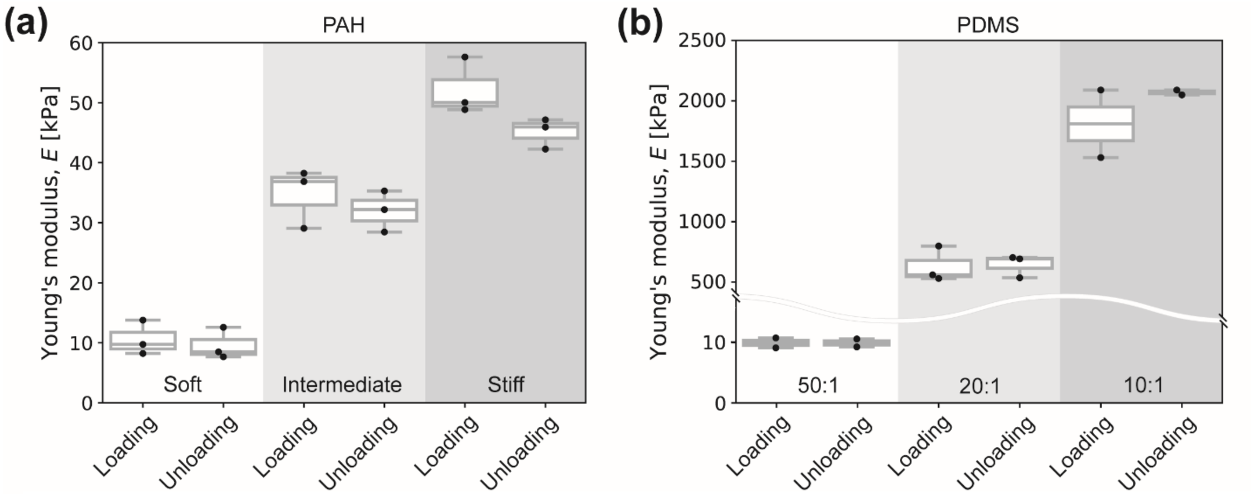


**SI Figure 1.** Young's modulus values comparison obtained from loading and unloading to confirm elasticity for (a) PAH and (b) PDMS.

We extracted *E* during loading and unloading dead weights to confirm that the reported Young's modulus values were within the elastic (reversible) range, as shown in SI Figure 1. The numerical values are summarized in SI Table 4.

**SI Table 4.** Average of loading and unloading for both PAH and PDMS

| **Sample** | **Young's modulus, $E_{Loading}$ (kPa)** | **Young's modulus, $E_{Unloading}$ (kPa)** |
|---|---|---|
| Soft PAH | 10.58 ± 2.87 | 9.58 ± 2.63 |
| Intermediate PAH | 34.72 ± 4.95 | 31.98 ± 3.43 |
| Stiff PAH | 52.16 ± 4.76 | 45.11 ± 2.54 |
| 50:1 PDMS | 9.93 ± 1.16 | 9.91 ± 0.93 |
| 20:1 PDMS | 622.37 ± 147.07 | 636.61 ± 93.61 |
| 10:1 PDMS | 1802.48 ± 395.17 | 2061.86 ± 28.31 |

**Effects of pre-strain on bulk rheology measurements**

To evaluate sample response under compression of stiff PAH and 10:1 PDMS, we conducted bulk shear rheology measurements under 1%, 2%, and 3% pre-compression strain. SI Figure 2(a) and (b) shows the log-log curves of $G'$ for stiff PAH as a function of $\gamma$ and $\omega$, respectively. These results confirm that all PAH bulk rheological responses did not significantly change with increasing compression for small shear strains (i.e., 0.01-10%). The initial average $G'$ values measured from the shear strain sweeps were 10.25 ± 1.83 kPa, 8.14 ± 0.55 kPa, and 9.60 ± 3.52 kPa, for 1%, 2%, and 3% compression respectively. The initial average $G'$ values measured from frequency sweep measurements were 10.7 ± 1.0 kPa, 9.80 ± 0.25 kPa, and 9.96 ± 0.2 kPa for 1%, 2%, and 3%, respectively.

PDMS revealed increasing storage modulus as compression increased for small shear strains (*i.e.*, 0.01-10%). The initial average $G'$ values measured from the shear strain sweeps were 67.93 ± 21.4 kPa, 84.91 ± 14.5 kPa, 93.45 ± 4.30 kPa for 1%, 2%, and 3% compression, respectively. The initial average $G'$ values measured from frequency sweep measurements were 56.1 ± 14.7 kPa, 64.3 ± 6.8 kPa, 80.73 ± 0.99 kPa for 1%, 2%, and 3%, respectively.

**Table 5.** Shear moduli obtained from bulk rheology with various pre-compression strains.

| Sample | Compression | Shear modulus, $G'_{StrainSweep}$ (kPa) | Shear modulus, $G'_{FrequencySweep}$ (kPa) |
|---|---|---|---|
| Stiff PAH | 1% compression | 10.25 ± 1.83 | 10.7 ± 1.0 |
| | 2% compression | 8.14 ± 0.55 | 9.80 ± 0.25 |
| | 3% compression | 9.60 ± 3.52 | 9.96 ± 0.2 |
| 10:1 PDMS | 1% compression | 67.93 ± 21.4 | 56.1 ± 14.7 |
| | 2% compression | 84.91 ± 14.5 | 64.3 ± 6.8 |
| | 3% compression | 93.45 ± 4.30 | 80.73 ± 0.99 |

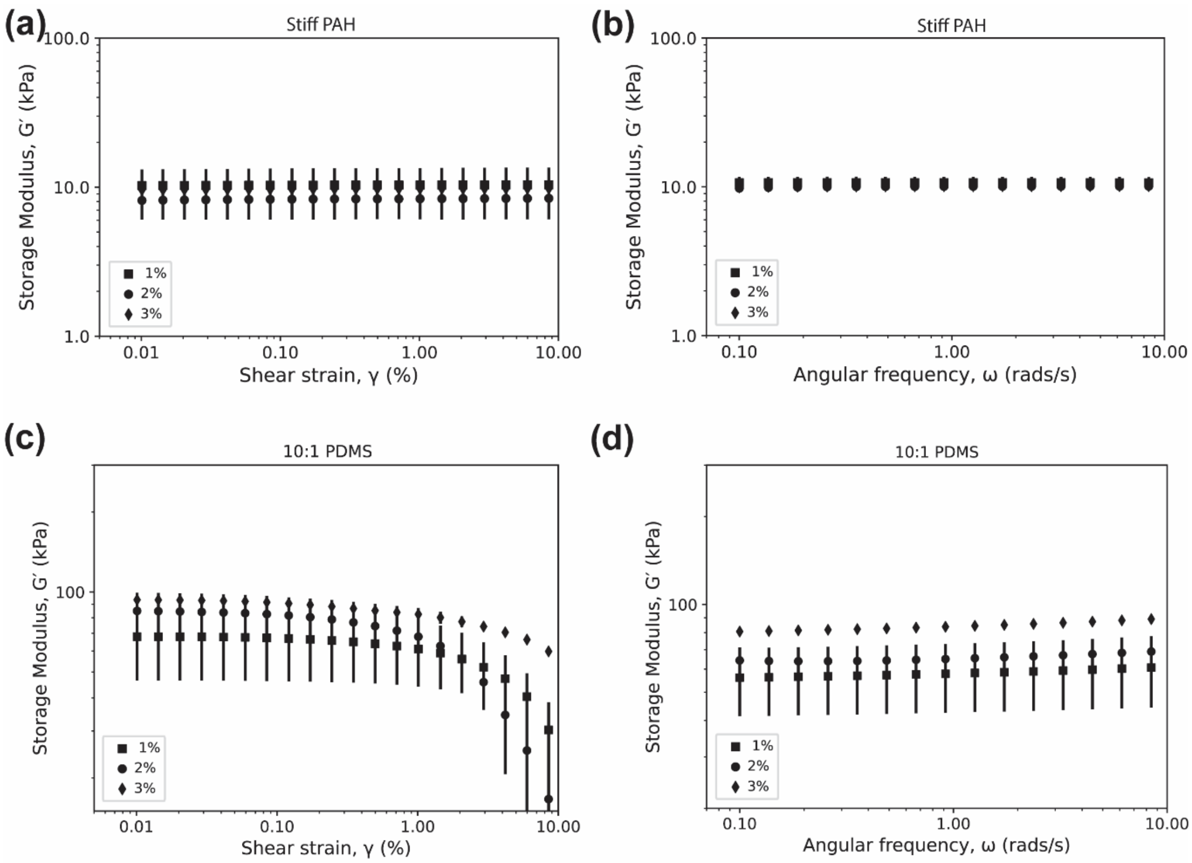


**SI Figure 2.** Effect of pre-compression strain on the bulk rheological response of stiff PAH as a function of (a) shear strain and (b) angular frequency. Effect of pre-compression strain on the bulk rheological response of 10:1 PDMS as a function of (c) shear strain and (d) angular frequency.

### Swelling of PAH

To determine if the PAH samples polymerized between the parallel plates of the shear rheometer were fully swollen, we quantified the water uptake after polymerization after 12 hrs and 24 hrs fully immersed in milliQ water. Soft, intermediate, and stiff PAH resulted in a swelling ratio of 3.1 ± 0.6%, 16.4 ± 0.9%, and 1.7 ± 0.1% indicating the sample is very close to fully swollen directly after polymerization. After 24 hours of the sample being in miliQ water the swelling ratio was measure, the soft, intermediate, and stiff PAH sample water intake was 0.5%, 5%, and 0%, more than measured at 12 hours after polymerization as shown in SI Figure 3.

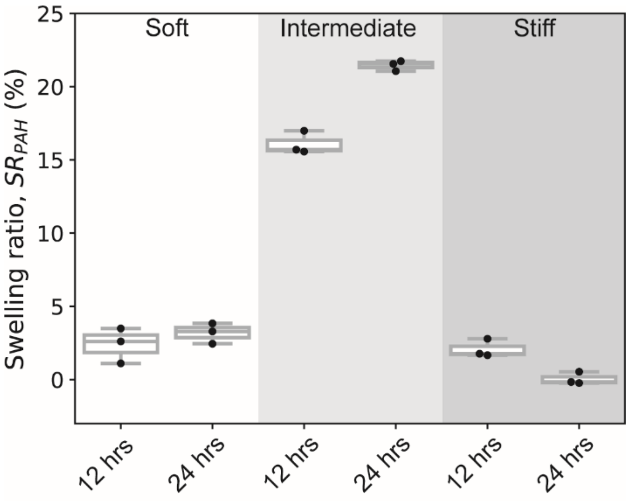


**SI Figure 3.** Swelling ratio obtained after 12 and 24 hrs for soft, intermediate, and stiff PAH compared against the as cast condition. Note that the as cast conditions already contains a significant amount of water.

**SI Table 6.** Swelling ratios for soft, intermediate, and stiff PAH after 12 hrs and 24 hrs immersed in milliQ water.

| **Sample** | **SR** $_{\mathbf{average}}$**, @ 12 hr (%)** | **SR** $_{\mathbf{average}}$**, @ 24 hr (%)** |
|---|---|---|
| Soft PAH | 2.41 ± 1.2 | 3.2 ± 0.70 |
| Intermediate PAH | 16.1 ± 0.78 | 21.4 ± 0.34 |
| Stiff PAH | 2.08 ± 0.62 | 0.05 ± 0.43 |

**SI Table 7.** Estimated mesh sizes obtained from $G_{\mathrm{StrainSweep}}$ and $G_{\mathrm{FrequencySweep}}$ using Flory's and de Gennes theories.

| **Sample** | **Shear modulus,** $G_{\mathbf{average}}$ **(kPa)** | **Mesh size, ξ (nm)** |
|---|---|---|
| Soft PAH | 2.4 ± 0.5 | 12 |
| Intermediate PAH | 6.9 ± 0.5 | 8 |
| Stiff PAH | 10.5 ± 0.5 | 7 |
